# A Deep Learning solution for triaging patient with cancer from their predicted mutational status using histopathological images

**DOI:** 10.1101/2022.01.30.475484

**Authors:** Louis-Oscar Morel, Valentin Derangère, Laurent Arnould, Sylvain Ladoire, Nathan Vinçon

## Abstract

The presence of mutation in cancer can be associated with a response to a targeted cancer therapy. Therefore it has become an important information while it helps giving a more specialized and more efficient treatment for each patient. Detection of mutation is routinely made by DNA-sequencing diagnostic tests. Recently, a new strategy of deep learning based method of prediction of mutation from histopathological images has shown promising results. However it is still unknown whether these methods can be useful as screening tests aside from sequencing methods for efficient population diagnosis. Here we developed our own deep learning based prediction pipeline for the detection of mutation in breast, lung, colon and ovarian cancer and find clinically relevant genes that have easy-to-detect mutational status, and especially *TP53* which can be detected in these four tumors. We then propose 3 potential screening strategies and show the effectiveness of our deep learning pipeline for optimizing diagnosis in the patient population.

## Introduction

Targeted cancer therapies are specialized and efficient therapies that have revolutionized the treatment of cancer in the last few years (1, 2). The higher specialization of targeted cancer therapies requires to know more and more information about the patient and getting personalized information requires making more specialized diagnostic tests (3). As an example, the presence or the absence of genomic mutations can be associated with a response to a targeted cancer therapy like PARPs (4) or WEE1 (5) inhibitors are treatments that are efficient only on cancers where *BRCA1/2* or *TP53* are mutated respectively. Detection of somatic mutation is routinely made by DNA-sequencing. However, these tests face a three fold limitation: they have a long waiting period, require a large amount of tissue and are expensive. Therefore, there is a growing need to identify new biomarkers and associated screening strategies to improve diagnostic workflows efficiency in medical oncology.

Recently, deep learning methods have shown promising results for the prediction of the mutational status from digitized tissue stained with hematoxylin and eosin as whole slide images (WSI) (6–12). These WSI are already made routinely in the diagnostic workflow and deep learning methods are cheap, always feasible and very scalable. Therefore, a deep learning based solution assessing the patient’s tumoral mutational status directly onto the WSI appears to be a potential screening strategy.

Here, we develop a deep learning pipeline to predict mutational status directly from WSI and find multiple predictable mutated genes. We then create 3 screening strategies to test the relevance of the deep learning pipeline in a routine context. The first screening strategy is “Save-all” which considers the number of diagnostic tests we can avoid while preserving high sensitivity. The second strategy “Fixed-Capacity” which considers, in the case of a limited number of diagnostic tests, the number of mutated patients found. In other words, it optimizes the number of patients that will later benefit the associated targeted therapy. The last screening strategy is “Prioritization” which considers the number of mutated patients found in a small part of the patient population for short-tracking. The rationale behind this strategy is that the earlier the patient has access to the best therapy, the higher its chance of remission. We finally show the relevance of our deep learning algorithm for each strategy in different realistic screening scenarii by showing its efficiency for each gene that both has a predictable mutational status and is relevant because it has an associated targeted therapy on the market or in development. A previous use of deep learning in a screening strategy were described in Nielsen *et al*. (13), but to our knowledge, this is the first screening strategy described for the prediction of mutational status in cancer.

## Material and Methods

### Study Design

All experiments were conducted in accordance with the Declaration of Helsinki and the International Ethical Guidelines for Biomedical Research Involving Human Subjects. Anonymized scanned WSIs were retrieved from the TCGA project through the Genomic Data Commons Portal (https://portal.gdc.cancer.gov/). We applied our method to the following tumor types: breast (BRCA), lung (LUAD), colon (COAD) and ovary (OV).

Ethics oversight of the TCGA study is described at https://www.cancer.gov/about-nci/organization/ccg/research/structural-genomics/tcga/history/policies. Informed consent was obtained by all participants in the TCGA.

### Datasets

All data, including histological images and information about the participants from the TCGA database are available at https://portal.gdc.cancer.gov/. Genetic data for patients in the TCGA cohorts are available at https://portal.gdc.cancer.gov/. The corresponding authors of this study are not involved in data sharing decisions of the TCGA database.

#### Molecular labels

Molecular labels were determined from the masked somatic mutations maf file of somatic mutation using the MuTect2 (14) algorithm corresponding to the dataset. An IMPACT value of “HIGH” or “MODERATE” categorized by VEP software (15) was considered positive while other values were considered negative.

### Pipeline

All analyses were performed using Python.

#### Image preprocessing

Aperio SVS files of diagnostic slides (labeled by DX in their name) from the 4 datasets were first selected. We then extracted the foreground, then tiled the images in non-overlapping patches of 600×600 pixels. These patches are associated with the label 1 if the gene is muted, 0 if the gene is non muted. These patches and labels were used as inputs of the EfficientNetB7 (16) neural network. Output prediction was a float between 0 and 1 for each patch. For the slide-level scoring, we extracted the 99th percentile of the patch prediction score distribution.

#### Neural network training, model selection

We used an EfficientNetB7 neural network architecture with a single sigmoid output layer after the global average pooling layer. The model was trained for 5 epochs where each epoch included all patches without further refinement.

### Implementation and hardware

Experiments were run with a NVIDIA RTX A4000 graphic card and TensorFlow v2.8.0-rc0, keras v2.8.0, CUDA 11.5.

## Results

### A significant proportion of mutations are detectable with deep learning methods on whole-slide imaging

We set up a deep learning pipeline predicting solid tumor mutations from WSI based on an EfficientNetB7 (16) pretrained on ImageNet dataset. In this study, 4 datasets from GDC Portal were analyzed, a lung cancer dataset, a colorectal cancer dataset, a breast cancer dataset and an ovarian cancer dataset, respectively TCGA-LUAD, TCGA-COAD, TCGA-BRCA and TCGA-OV. We systematically trained the deep learning pipeline on every mutation having a prevalence greater than 10% in the dataset to focus on mutations that can show statistical significance. These correspond to 922 tested genes. On TCGA-LUAD, TCGA-COAD and TCGA-BRCA, 43% of the gene had a statistically significant predictable mutational status (Figure 1). The TCGA-OV dataset had too few cases to reach statistical significance for most of the genes, therefore we removed it for later analyses. However, we noticed that the *TP53* gene had a predictable mutational status with an area under the roc curve (AUC) of 0.65.

**Fig. 1.**
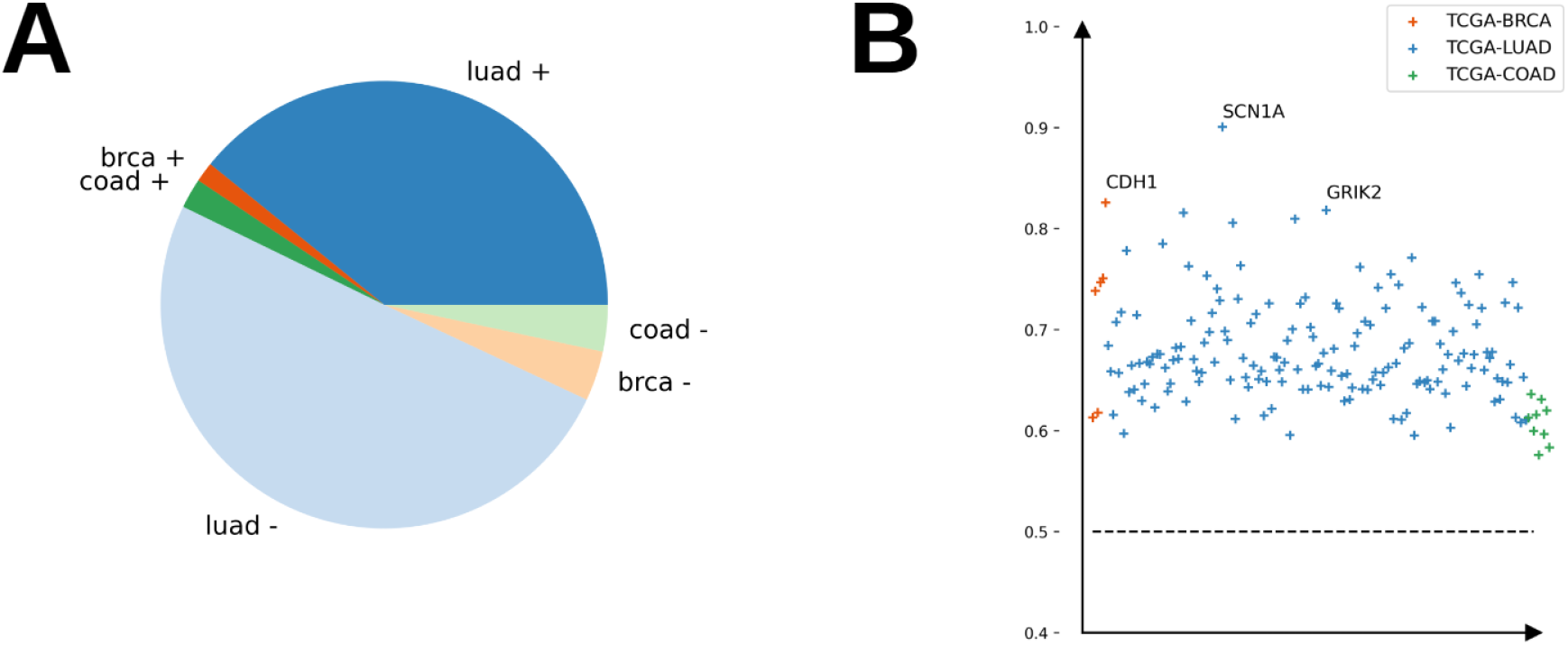
Description of predictable genes in TCGA-LUAD, TCGA-COAD and TCGA-BRCA. Pie of the number of genes according to their dataset and their predictability (see Material and Methods). Predictable genes are shown in dark with “+”, other genes are shown in light with “-”. Scatter plot of the AUC of predictable genes. The dashed bar at 0.5 corresponds to the AUC of a random classifier. Dataset of origin is shown in color.

### Mutations on targetable genes are detectable

Among the most detectable mutations, we found *SCN1A* in TCGA-LUAD with an AUC of 0.90, *CDH1* in TCGA-BRCA with an AUC of 0.83 and *GRIK2* in TCGA-LUAD with an AUC of 0.82 (Figure 1). These results show a high morphological effect of many types of molecular alterations. We then look at genes that were relevant as clinical information because they are associated with either a clinically approved drug or an in-development drug. For convenience, we later call the former not clinically relevant genes and the latter clinically relevant genes respectively non-clinical genes and clinical genes. In clinical genes, we found that mutations in *TP53* were detectable in TCGA-COAD, TCGA-LUAD, TCGA-BRCA and TCGA-OVARY with an AUC of 0.60, 0.64, 0.74 and 0.65 respectively (Table 1). *TP53* was the only gene found detectable in the 4 datasets which suggest that *TP53* is ubiquitously detectable and could have a pan-tumoral morphological signature; *TP53* was significantly easier to find in the TCGA-BRCA dataset. In addition, *KRAS* mutations were found detectable in both TCGA-COAD and TCGA-LUAD with an AUC of 0.63 and 0.65 respectively. Also, mutations in the *APC* gene were detectable in TCGA-COAD with an AUC of 0.61, *EGFR* mutations were detectable in TCGA-LUAD with an AUC of 0.61. These results are similar to the pan-cancer study in (10), thus confirming their results.

**Table 1.**
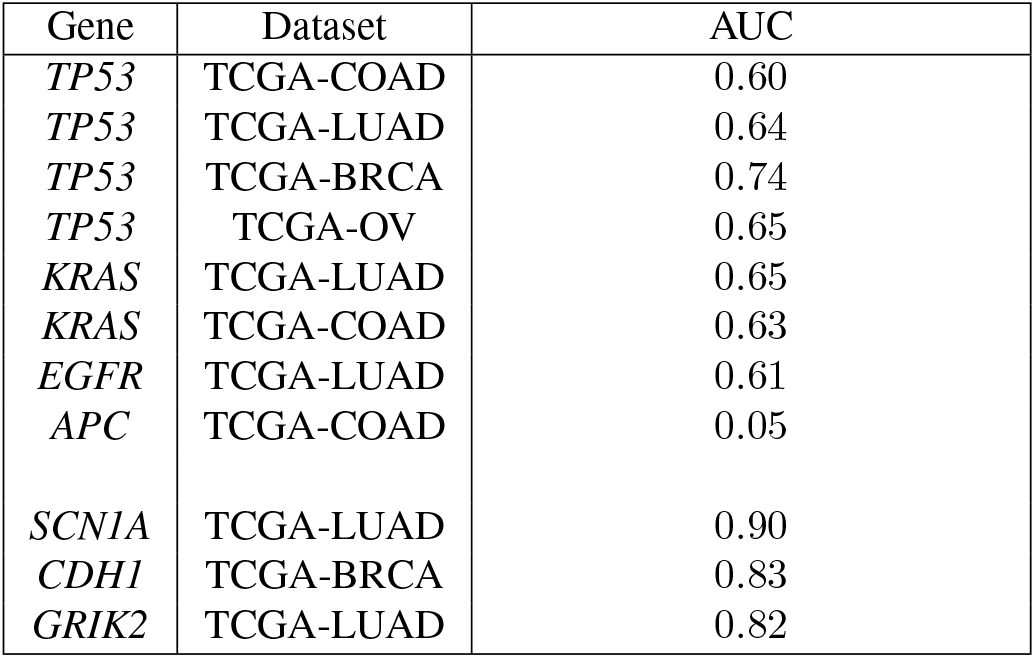
AUC scores for clinically relevant (top) and non clinically relevant (bottom) genes.

| Gene | Dataset | AUC |
| --- | --- | --- |
| *TP53* | TCGA-COAD | 0.60 |
| *TP53* | TCGA-LUAD | 0.64 |
| *TP53* | TCGA-BRCA | 0.74 |
| *TP53* | TCGA-OV | 0.65 |
| *KRAS* | TCGA-LUAD | 0.65 |
| *KRAS* | TCGA-COAD | 0.63 |
| *EGFR* | TCGA-LUAD | 0.61 |
| *APC* | TCGA-COAD | 0.05 |
| *SCN1A* | TCGA-LUAD | 0.90 |
| *CDH1* | TCGA-BRCA | 0.83 |
| *GRIK2* | TCGA-LUAD | 0.82 |

### Screening strategies can optimize the diagnosis in the patient population

In the next step, we analyze the performance of the most predictable of the previously identified genes in realistic screening scenarii. We define 3 potential strategies and show the associated performance of the algorithm as a discriminative test. First strategy is “Save-all”, where we ask whether the algorithm can filter non-mutated patients while having almost no false-negative in order to avoid unnecessary diagnostic tests. More specifically, we are interested in the number of diagnostic tests that can be avoided while preserving a sensitivity of 95% or 99% in our screening test (Figure 2). Second strategy is “Fixed-Capacity”, where we ask whether we can optimize the number of mutated patients found in the case of a limited number of sequencing tests. In this configuration, the goal is to optimize the number of patients that will finally benefit from the associated targeted therapy. More specifically, we are interested in the increase of the NGS sensitivity if their capacity to test is 30%, 50% or 70% of the patient population (Figure 2). Third strategy is “Prioritize” where we ask whether we can find a small proportion of patients that are highly likely to be positive in order to prioritize them for the sequencing diagnostic test. The rationale behind the “Prioritize” strategy is that in a context of cancer, time is against the patient and days can significantly change the prognosis (17). More specifically, we are interested in the positive predictive value (PPV) for a cutoff of the 5% and 10% most likely to be mutated for the given gene (Figure 2). We have chosen 5% and 10% cutoffs such that it is high enough to have an influence at the population level and low enough to be realistic for a short-track.

**Fig. 2.**
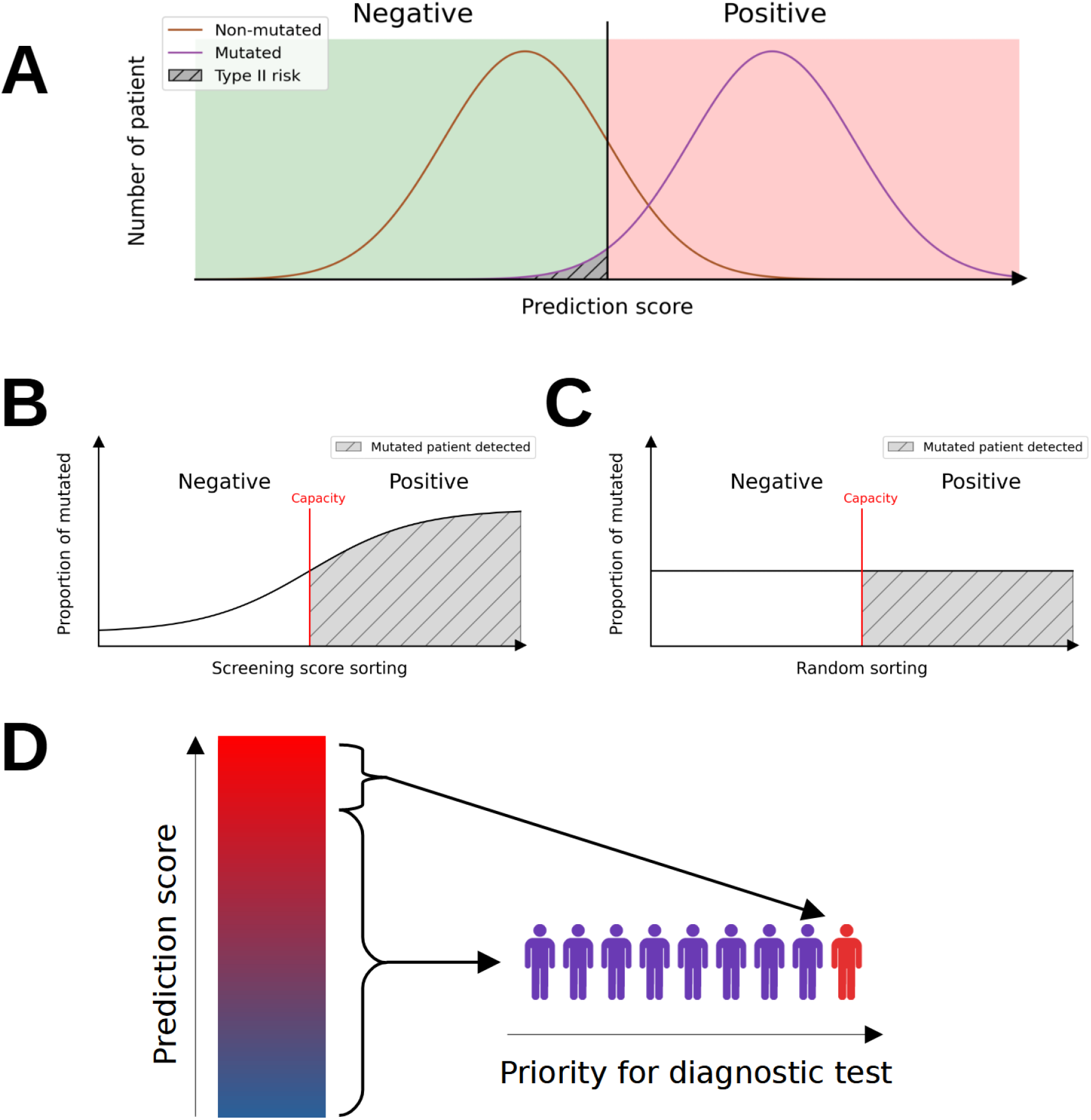
Screening strategies of the deep learning triage pipeline. (A) Save-all strategy, violet and brown gaussian curves are respectively the number of patients with and without a given mutation. The red zone corresponds to the positive patient for the screening test and green zone the negative patient. The gray area is the type II error (false negative) taking the value of 5% and 1% in our scenarii. (B) and (C) Fixed-capacity strategy respectively using the screening test and using random test. The red vertical line is the limit capacity of diagnostic tests, therefore all patients having a higher score than the limit capacity are diagnosed. The y-axis corresponds to the proportion of mutated patients for a given screening score. The screening strategy is expected to find more mutated patients than the random strategy using the same number of diagnosis tests. (D) Priority strategy, the blue-to-red gradient corresponds to the screening score, higher score is in red. The top 5% or 10% patients are selected for priority diagnostic tests because they are very likely to be positive.

For each screening strategy our deep learning pipeline performance is compared to a random screening test. Intuitively, the random screening test would provide 5% and 1% avoided tests for 95% and 99% sensitivity respectively.

In the “Save-all” strategy (Figure 3), for the non-clinical genes, proportion of avoided tests reached 73% for *SCN1A* in TCGA-LUAD dataset (Figure 3). For the clinical genes, the proportion of avoided tests using screening range from 7.8 to 26.3 for a 95% sensitivity and range from 2.6 to 14.1 for a 99% sensitivity. This means that a quarter of the diagnostic tests can be avoided in the best case, which is with the *EGFR* gene in TCGA-LUAD. For 95% sensitivity, in all genes but *APC* in TCGA-COAD, more than 10% tests could be avoided and other genes with highest score were *KRAS* in TCGA-LUAD with 14.4%, *TP53* in TCGA-LUAD with 14.7% and TP53 in TCGA-BRCA with 18.5%.

**Fig. 3.**
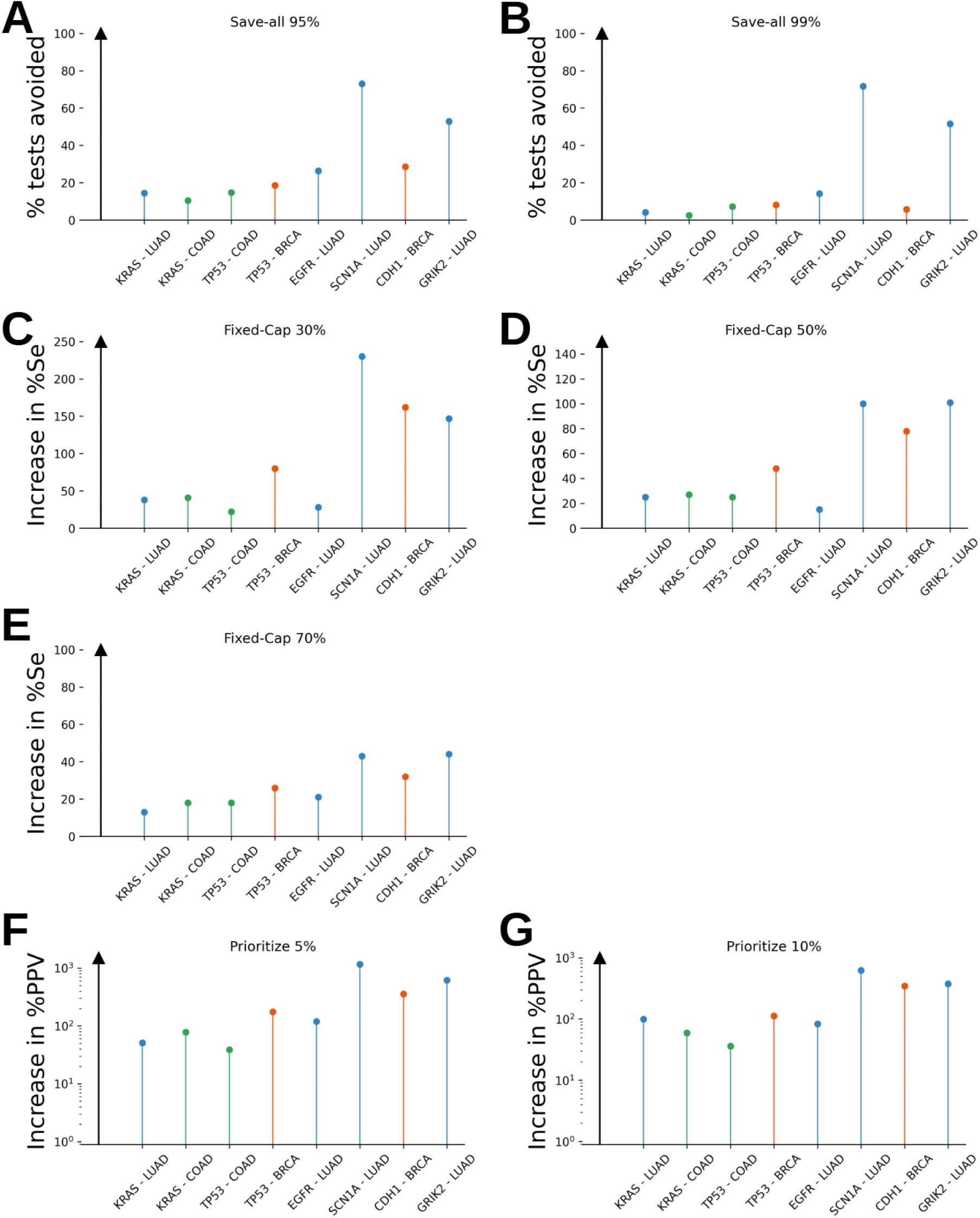
Performance of the screening pipeline for the 3 screening strategies on multiple clinical and non-clinical genes. (A) and (B) Percentage of avoided tests at a sensitivity threshold of 5% and 1% respectively (save-all strategy). The colors correspond to the dataset of origin using the same code as in figure 1. (C-E) Percentage increase in sensitivity for the fixed-capacity strategy with a capacity of respectively 30%, 50% and 70%. The colors correspond to the dataset of origin using the same code as in figure 1. (FG) Percentage increase in positive predictive value for the priority strategy with a threshold of 5% and 10%. The y-axis is shown in log scale. The colors correspond to the dataset of origin using the same code as in figure 1.

In the “Fixed-Capacity” strategy (Figure 3), for the non-clinical genes, sensitivity improvement reached +230% for *SCN1A* in TCGA-LUAD for a capacity of 30% and +101% for *GRIK2* in TCGA-LUAD for a capacity of 50%. For the clinical genes, the sensitivity improvement ranges from +19% to +80% for a capacity of 30% of the patient population, from +15% to +48% for a capacity of 50% and from +13% to +26% for a capacity of 70%. Intuitively, the lower is the capacity, the higher is the beneficial effect of the screening strategy. For a 30% capacity, the genes with highest score were *TP53* in TCGA-BRCA with +80% sensitivity improvement, then *KRAS* in TCGA-COAD with +41% and *KRAS* in TCGA-LUAD with +38% improvement.

In the “Prioritize” strategy (Figure 3), for the non-clinical genes, the increase in PPV reached +1166% for *SCN1A* in TCGA-LUAD with a 5% cutoff and +633% for the same gene with a 10% cutoff. For the clinical genes, the increase in PPV ranges from +24% to +181% with a 5% cutoff and from +12% to +117% with a 10% cutoff. Best performances were once more obtained by *TP53* in TCGA-BRCA dataset, and *EGFR* in TCGA-LUAD have an increase in PPV of +125% (2.25 times higher risk of being mutated when our test is positive) and +87% for a respective cutoff of 5% and 10%.

Interestingly, having a high AUC score is not necessarily associated with having better performance in the different screening scenarii, the difference can mostly be explained by the prevalence of the mutation in the patient population. Intuitively, the less there are mutated patients, the more the screening can help find more of them with a higher magnitude.

In conclusion, among the 3 proposed strategies, the “Fixed-Capacity strategy” shows that screening can provide significant improvement in the case where only a limited number of diagnostic tests can be done. The “Prioritize strategy” appears to be efficient for a 5% cutoff but quickly becomes less efficient at a 10% cutoff. Finally, while the “Save-all strategy” can drastically reduce the number of patients to test for non-clinical genes, its effect for clinical genes is less important as it only removes 18.5% of the population for a sensitivity of 95% in the best case. These results would require a medico-economic analysis to clearly see potential advantages of a deep-learning based screening in application.

## Discussion

Cancer is a very diversified disease where no treatment is ubiquitously sufficiently efficient. Targeted therapies have emerged in response to this diversity, being specialized on subtypes of cancer and belonging to a more personalized medicine. However these new treatments require acquiring more information about the cancer by characterizing its molecular and mutational statuses. Acquiring such information can be very expensive, especially for innovative DNA-sequencing methods which, added to their lack of scalability, makes their democratization difficult. In this paper, we propose credible screening strategies based on new deep learning methods, cheaper, quicker and more scalable. We show that predicting mutational status from WSI is efficient and can optimize the allocation of diagnostic test resources in 3 credible scenarii for clinically relevant genes *TP53*, *KRAS* and *EGFR*.

Clinically relevant genes were defined as genes having an approved or in-development drug associated with their mutational status. *TP53* mutation is an indication for the WEE1 inhibitor in a clinical trial of phase 3 in pancreas cancer, for ibrutinib (18) and idelalisib (19) in chronic lymphocytic leukemia. The absence of *KRAS* mutations is an indication for cetuximab (20) in colorectal cancer. Finally, *EGFR* mutation is an indication for erlotinib (21) or gefitinib (22) in lung cancer.

As for each cancer type only one dataset was used, we cannot provide an estimation of the robustness of mutation detectability to change in the data distribution that would be caused by images coming from a new center. In other words, robustness to inter-center variation is completely unknown for the specific task of mutational status prediction. This is a known barrier for clinical application of deep learning based methods. In this paper, we approximated the population prevalence by the dataset prevalence for the mutation. While prevalence does not affect sensitivity, it has influence over the PPV and the threshold of a given sensitivity. Therefore, results shown for “Fixed-Capacity” strategy would be similar for small prevalence adjustment but results for “Prioritize” and “Save-all” strategy are highly dependent on the true mutation prevalence.

To describe benefits of the shown screening strategy more accurately, a medico-economic analysis could be done integrating the cost and availability of a diagnostic test in a first part, then the clinical benefits of the considered targeted therapy in a second part. Finally, with the ever-changing possibilities in medical cancerology, non-clinically relevant genes but with highly detectable mutational status might become relevant in the future.

## Bibliography

1. Kelly E. McCann, Sara A. Hurvitz, and Nicholas McAndrew. Advances in Targeted Therapies for Triple-Negative Breast Cancer. Drugs, 79(11):1217–1230, July 2019. ISSN 1179-1950. doi: 10.1007/s40265-019-01155-4.

2. Sabine Kayser and Mark J. Levis. Advances in targeted therapy for acute myeloid leukaemia. British Journal of Haematology, 180(4):484–500, 2018. ISSN 1365-2141. doi: 10.1111/bjh.15032. _eprint: https://onlinelibrary.wiley.com/doi/pdf/10.1111/bjh.15032.

3. Meagan B Myers. Targeted therapies with companion diagnostics in the management of breast cancer: current perspectives. Pharmacogenomics and Personalized Medicine, 9: 7–16, January 2016. ISSN 1178-7066. doi: 10.2147/PGPM.S56055.

4. Karolina N. Dziadkowiec, Emilia Gąsiorowska, Ewa Nowak-Markwitz, and Anna Jankowska. PARP inhibitors: review of mechanisms of action and BRCA1/2 mutation targeting. Przeglad Menopauzalny = Menopause Review, 15(4):215–219, December 2016. ISSN 1643-8876. doi: 10.5114/pm.2016.65667.

5. Suzanne Leijen, Robin M.J.M. van Geel, Gabe S. Sonke, Daphne de Jong, Efraim H. Rosenberg, Serena Marchetti, Dick Pluim, Erik van Werkhoven, Shelonitda Rose, Mark A. Lee, Tomoko Freshwater, Jos H. Beijnen, and Jan H.M. Schellens. Phase II Study of WEE1 Inhibitor AZD1775 Plus Carboplatin in Patients With *TP53*-Mutated Ovarian Cancer Refractory or Resistant to First-Line Therapy Within 3 Months. Journal of Clinical Oncology, 34(36):4354–4361, December 2016. ISSN 0732-183X, 1527-7755. doi: 10.1200/JCO.2016.67.5942.

6. Nicolas Coudray, Paolo Santiago Ocampo, Theodore Sakellaropoulos, Navneet Narula, Matija Snuderl, David Fenyö, Andre L. Moreira, Narges Razavian, and Aristotelis Tsirigos. Classification and mutation prediction from non–small cell lung cancer histopathology images using deep learning. Nature Medicine, 24(10):1559–1567, October 2018. ISSN 1546-170X. doi: 10.1038/s41591-018-0177-5. Number: 10 Publisher: Nature Publishing Group.

7. Pierre Murchan, Cathal Ó’Brien, Shane O’Connell, Ciara S. McNevin, Anne-Marie Baird, Orla Sheils, Pilib Ó Broin, and Stephen P. Finn. Deep Learning of Histopathological Features for the Prediction of Tumour Molecular Genetics. Diagnostics, 11(8):1406, August 2021. ISSN 2075-4418. doi: 10.3390/diagnostics11081406. Number: 8 Publisher: Multidisciplinary Digital Publishing Institute.

8. Mohsin Bilal, Shan E Ahmed Raza, Ayesha Azam, Simon Graham, Mohammad Ilyas, Ian A Cree, David Snead, Fayyaz Minhas, and Nasir M Rajpoot. Development and validation of a weakly supervised deep learning framework to predict the status of molecular pathways and key mutations in colorectal cancer from routine histology images: a retrospective study. The Lancet Digital Health, 3(12):e763–e772, December 2021. ISSN 2589-7500. doi: 10.1016/S2589-7500(21)00180-1.

9. Sidong Liu, Zubair Shah, Aydin Sav, Carlo Russo, Shlomo Berkovsky, Yi Qian, Enrico Coiera, and Antonio Di Ieva. Isocitrate dehydrogenase (IDH) status prediction in histopathology images of gliomas using deep learning. Scientific Reports, 10(1):7733, May 2020. ISSN 2045-2322. doi: 10.1038/s41598-020-64588-y. Number: 1 Publisher: Nature Publishing Group.

10. Javad Noorbakhsh, Saman Farahmand, Mohammad Soltanieh-ha, Sandeep Namburi, Kourosh Zarringhalam, and Jeff Chuang. Pan-cancer classifications of tumor histological images using deep learning. Technical report, bioRxiv, July 2019. Section: New Results.

11. Hyun-Jong Jang, Ahwon Lee, J Kang, In Hye Song, and Sung Hak Lee. Prediction of clinically actionable genetic alterations from colorectal cancer histopathology images using deep learning. World Journal of Gastroenterology, 26(40):6207–6223, October 2020. ISSN 1007-9327. doi: 10.3748/wjg.v26.i40.6207.

12. Hyun-Jong Jang, Ahwon Lee, Jun Kang, In Hye Song, and Sung Hak Lee. Prediction of genetic alterations from gastric cancer histopathology images using a fully automated deep learning approach. World Journal of Gastroenterology, 27(44):7687–7704, November 2021. ISSN 1007-9327. doi: 10.3748/wjg.v27.i44.7687.

13. Katrine B. Nielsen, Mie L. Lautrup, Jakob K. H. Andersen, Thiusius R. Savarimuthu, and Jakob Grauslund. Deep Learning–Based Algorithms in Screening of Diabetic Retinopathy: A Systematic Review of Diagnostic Performance. Ophthalmology Retina, 3(4):294–304, April 2019. ISSN 2468-6530. doi: 10.1016/j.oret.2018.10.014.

14. Kristian Cibulskis, Michael S. Lawrence, Scott L. Carter, Andrey Sivachenko, David Jaffe, Carrie Sougnez, Stacey Gabriel, Matthew Meyerson, Eric S. Lander, and Gad Getz. Sensitive detection of somatic point mutations in impure and heterogeneous cancer samples. Nature Biotechnology, 31(3):213–219, March 2013. ISSN 1546-1696. doi: 10.1038/nbt.2514.

15. William McLaren, Laurent Gil, Sarah E. Hunt, Harpreet Singh Riat, Graham R. S. Ritchie, Anja Thormann, Paul Flicek, and Fiona Cunningham. The Ensembl Variant Effect Predictor. Genome Biology, 17(1):122, June 2016. ISSN 1474-760X. doi: 10.1186/s13059-016-0974-4.

16. Mingxing Tan and Quoc V. Le. EfficientNet: Rethinking Model Scaling for Convolutional Neural Networks. arXiv:1905.11946 [cs, stat], September 2020. arXiv: 1905.11946.

17. Louise Eriksson, Jonas Bergh, Keith Humphreys, Fredrik Wärnberg, Sven Törnberg, and Kamila Czene. Time from breast cancer diagnosis to therapeutic surgery and breast cancer prognosis: A population-based cohort study. International Journal of Cancer, 143(5):1093–1104, September 2018. ISSN 1097-0215. doi: 10.1002/ijc.31411.

18. Jan A. Burger, Paul M. Barr, Tadeusz Robak, Carolyn Owen, Paolo Ghia, Alessandra Tedeschi, Osnat Bairey, Peter Hillmen, Steven E. Coutre, Stephen Devereux, Sebastian Grosicki, Helen McCarthy, David Simpson, Fritz Offner, Carol Moreno, Sandra Dai, Indu Lal, James P. Dean, and Thomas J. Kipps. Long-term efficacy and safety of first-line ibrutinib treatment for patients with CLL/SLL: 5 years of follow-up from the phase 3 RESONATE-2 study. Leukemia, 34(3):787–798, March 2020. ISSN 1476-5551. doi: 10.1038/s41375-019-0602-x.

19. Jeffrey A Jones, Tadeusz Robak, Jennifer R Brown, Farrukh T Awan, Xavier Badoux, Steven Coutre, Javier Loscertales, Kerry Taylor, Elisabeth Vandenberghe, Malgorzata Wach, Nina Wagner-Johnston, Loic Ysebaert, Lyndah Dreiling, Ronald Dubowy, Guan Xing, Ian W Flinn, and Carolyn Owen. Efficacy and safety of idelalisib in combination with ofatumumab for previously treated chronic lymphocytic leukaemia: an open-label, randomised phase 3 trial. The Lancet Haematology, 4(3):e114–e126, March 2017. ISSN 2352-3026. doi: 10.1016/S2352-3026(17)30019-4.

20. Astrid Lièvre, Jean-Baptiste Bachet, Delphine Le Corre, Valérie Boige, Bruno Landi, Jean-François Emile, Jean-François Côté, Gorana Tomasic, Christophe Penna, Michel Ducreux, Philippe Rougier, Frédérique Penault-Llorca, and Pierre Laurent-Puig. KRAS Mutation Status Is Predictive of Response to Cetuximab Therapy in Colorectal Cancer. Cancer Research, 66(8):3992–3995, April 2006. ISSN 0008-5472, 1538-7445. doi: 10.1158/0008-5472.CAN-06-0191. Publisher: American Association for Cancer Research Section: Priority Reports.

21. Rafael Rosell, Enric Carcereny, Radj Gervais, Alain Vergnenegre, Bartomeu Massuti, Enriqueta Felip, Ramon Palmero, Ramon Garcia-Gomez, Cinta Pallares, Jose Miguel Sanchez, Rut Porta, Manuel Cobo, Pilar Garrido, Flavia Longo, Teresa Moran, Amelia Insa, Filippo De Marinis, Romain Corre, Isabel Bover, Alfonso Illiano, Eric Dansin, Javier de Castro, Michele Milella, Noemi Reguart, Giuseppe Altavilla, Ulpiano Jimenez, Mariano Provencio, Miguel Angel Moreno, Josefa Terrasa, Jose Muñoz-Langa, Javier Valdivia, Dolores Isla, Manuel Domine, Olivier Molinier, Julien Mazieres, Nathalie Baize, Rosario Garcia-Campelo, Gilles Robinet, Delvys Rodriguez-Abreu, Guillermo Lopez-Vivanco, Vittorio Gebbia, Lioba Ferrera-Delgado, Pierre Bombaron, Reyes Bernabe, Alessandra Bearz, Angel Artal, Enrico Cortesi, Christian Rolfo, Maria Sanchez-Ronco, Ana Drozdowskyj, Cristina Queralt, Itziar de Aguirre, Jose Luis Ramirez, Jose Javier Sanchez, Miguel Angel Molina, Miquel Taron, and Luis Paz-Ares. Erlotinib versus standard chemotherapy as first-line treatment for European patients with advanced EGFR mutation-positive non-small-cell lung cancer (EURTAC): a multicentre, open-label, randomised phase 3 trial. The Lancet Oncology, 13(3):239–246, March 2012. ISSN 1470-2045. doi: 10.1016/S1470-2045(11)70393-X.

22. Jean-Charles Soria, Yi-Long Wu, Kazuhiko Nakagawa, Sang-We Kim, Jin-Ji Yang, Myung-Ju Ahn, Jie Wang, James Chih-Hsin Yang, You Lu, Shinji Atagi, Santiago Ponce, Dae Ho Lee, Yunpeng Liu, Kiyotaka Yoh, Jian-Ying Zhou, Xiaojin Shi, Alan Webster, Haiyi Jiang, and Tony S K Mok. Gefitinib plus chemotherapy versus placebo plus chemotherapy in EGFR-mutation-positive non-small-cell lung cancer after progression on first-line gefitinib (IMPRESS): a phase 3 randomised trial. The Lancet Oncology, 16(8):990–998, August 2015. ISSN 1470-2045. doi: 10.1016/S1470-2045(15)00121-7.

